# Decoding dynamic implicit and explicit representations of facial expressions of emotion from EEG

**DOI:** 10.1101/453654

**Authors:** Fraser W. Smith, Marie L Smith

**Author notes:** Correspondence should be addressed to FWS. School of Psychology EDU Building University of East Anglia Norwich Research Park Norwich, NR4 7TJ.

## Abstract

Faces transmit a wealth of important social signals. While previous studies have elucidated the network of cortical regions important for perception of facial expression, and the associated temporal components such as the P100, N170 and EPN, it is still unclear how task constraints may shape the representation of facial expression (or other face categories) in these networks. In the present experiment, we investigate the neural information available across time about two important face categories (expression and identity) when those categories are either perceived under explicit (e.g. decoding emotion when task is on emotion) or implicit task contexts (e.g. decoding emotion when task is on identity). Decoding of both face categories, across both task contexts, peaked in a 100-200ms time-window post-stimulus (across posterior electrodes). Peak decoding of expression, however, was not affected by task context whereas peak decoding of identity was significantly reduced under implicit processing conditions. In addition, errors in EEG decoding correlated with errors in behavioral categorization under explicit processing for both expression and identity, but only with implicit decoding of expression. Despite these differences, decoding time-courses and the spatial pattern of informative electrodes differed consistently for both tasks across explicit Vs implicit face processing. Finally our results show that information about both face identity *and* facial expression is available around the N170 time-window on lateral occipito-temporal sites. Taken together, these results reveal differences and commonalities in the processing of face categories under explicit Vs implicit task contexts and suggest that facial expressions are processed to a richer degree even under implicit processing conditions, consistent with prior work indicating the relative automaticity by which emotion is processed. Our work further demonstrates the utility in applying multivariate decoding analyses to EEG for revealing the dynamics of face perception.

## Introduction

Successful decoding of facial expressions of emotion is an important social skill in humans. Hence understanding the neural mechanisms underlying this feat is an important question and the focus of much research. Studies using functional imaging have revealed a network of regions both within and beyond the core face network (Haxby et al., 2000) that show enhanced activity for emotional Vs neutral face stimuli (e.g Vuilleumier et al., 2001; Engell and Haxby, 2007). More recently, studies have investigated whether facial expression categories lead to different patterns of brain activity within regions of the face network (see Wegrezyn et al, 2015; Zhang et al, 2016; Greening et al 2018). These studies revealed that particular expression categories can be differentiated in multiple face and emotion brain regions, including superior temporal sulcus (STS - Wegrezyn et al, 2015; Zhang et al, 2016; Greening et al 2018), amygdala (Wegrezyn et al, 2015; Zhang et al, 2016) but also in the fusiform gyrus (FG) and inferior occipital gyrus (IOG) in particular contexts (Wegrezyn et al 2015; see also Greening et al 2018).

Interestingly, two of the aforementioned studies (Wegrezyn et al., 2015; Zhang et al., 2016), used orthogonal tasks which did not require participants to focus on expression to solve the task: i.e. a gender categorization task (Wegrezyn et al) or a fixation task (Zhang et al.), while Greening et al (2018) used an explicit emotion categorization task. Previous activation-based fMRI studies have shown that the nature of the task performed while viewing facial expressions (i.e. an explicit or implicit expression focus) can have a significant influence on the resulting brain activity observed (esp. e.g. in the amygdala; Critchley et al., 2000; Hariri et al 1999; see also Fusar-Poli et al 2009). However, at present it is still unclear how task may shape the *representation* of important face categories, such as facial expression in visual cortex (see Petro et al 2013; Kay & Yeatman, 2017). In the present work, we explicitly address this question using the complementary time-sensitive neuroimaging technique of EEG.

It is well known that neurons in the visual system, even extending back to primary visual cortex, receive significant amounts of top down influence via cortical feedback connections from multiple higher brain regions. It has been argued that these top down influences can effectively change the information conveyed by neurons by, for example, altering their response tuning (Gilbert & Li, 2013). The perceptual task an observer is asked to perform is one such top down influence that can change the activity of neurons in the visual system – even in V1 – to identical visual stimuli (Li et al 2004; Kay & Yeatman, 2017). This in turn allows for boosting of sensory representations that have behavioral relevance in a particular task context (Desimone & Duncan, 1995; see also Peelen et al 2009). While some evidence of task demands altering the response of early visual areas (Li et al 2004; Petro et al 2013) and higher visual areas (Kay & Yeatman, 2017) to identical visual stimuli exist, it is unclear to date how task may shape the dynamically evolving neural representation of important face categories such as expression (and identity).

While studies using fMRI have provided much valuable information about which brain regions code facial expressions, they are necessarily limited in their ability to speak to the time course of neural processing of facial expressions. Much research has been carried out using EEG and MEG, however, to address this question (see e.g. Eimer, 2011). Traditional ERP analyses have demonstrated a specific component related to face processing, the N170, that is thought to be generated in regions of the core face network, i.e. IOG, FG, STS or some combination thereof (see Eimer, 2011). Although the face sensitivity of the N170 is not in doubt, it is still unclear to what extent this component is sensitive to facial expression. Some studies did not report any sensitivity to emotion in the N170 (e.g. Eimer & Holmes, 2002; Eimer & Holmes, 2007; Pourtois et al 2005; Rellecke, Sommer & Schact, 2013; Neath-Tavares & Itier, 2016) and this is in keeping with a long-standing idea that the N170 indexes structural encoding of a face (see Eimer & Holmes, 2002). Other studies, however, do find some evidence of sensitivity to emotion on the N170 (Batty & Taylor, 2003; Leppanen et al., 2007, 2008; Schyns et al., 2007; M. Smith 2012; Turano et al 2017) and in fact a recent meta-analysis supports an effect of emotion for specific emotions (happy, angry and fear each > neutral) on the N170 (Hinojosa et al 2015). Recent evidence suggests that the earlier discrepancy could in part be driven by methodological differences including both the choice of reference electrodes (with linked mastoids dampening any such effect, Rellecke et al, 2013; Hinojosa et al 2015) and crucially, also by attentional focus (with indirect attention on expression leading to larger effects, Hinojosa et al 2015). In sum, at present it is still unclear to what extent the N170 component may carry discriminative information regarding *each* of the basic facial expressions (i.e. beyond emotion > neutral) across *both* explicit and implicit task contexts. In particular, within-subject designs directly comparing the effect of task on identical stimuli are ideally required to compellingly answer this question (Itier & Neath-Tavares, 2017). In the present work we test whether more sensitive multivariate analysis techniques (MVPA; see Grootswagers et al 2017, for a review) may provide novel evidence as to whether the EEG signal in the N170 time window permits the discrimination of the basic facial expressions of emotion (see Nemrodov et al., 2016, for a similar approach with respect to facial identity).

While the N170 has received arguably the most attention as a marker of face processing, visually-evoked components of interest typically begin around the P1, a positivity occurring 100ms post stimulus onset over extra-striate visual regions that is typically linked to low-level stimulus properties and attention (Rossion & Jacques, 2012; Luck et al, 2000). Although early emotion effects have been observed on the P1 in some studies (e.g. Batty & Taylor, 2003; Luo et al 2010; E. Smith et al, 2013), they are by no means consistently observed (see e.g. Itier & Neath-Tavares, 2017; Vulluimier & Pourtois, 2007) and do not necessarily indicate sensitivity to discrete emotions (e.g. though see Luo et al 2010). The early posterior negativity (EPN) component, occurring around 200-350ms is thought to reflect enhanced processing of emotion in extra-striate areas, likely reflecting feedback from higher brain areas (e.g. Itier & Neath-Tavares, 2017; Recio et al 2017; Pourtois et al 2012). This component is observed for emotional faces, words and pictures (see Itier & Neath-Tavares, 2017, for review) with higher amplitudes to emotional Vs neutral content and is therefore not thought to be face specific per se but rather linked to the encoding of emotional content in general. In particular some studies have found the EPN discriminates valence though not always in a consistent manner (see Itier & Neath-Tavares, 2017 for review). In addition, it been shown to be sensitive to task demands on identical facial expression stimuli (Itier & Neath-Tavares, 2017). These authors showed that differences between tasks were not evident until 200ms (on lateral occipito-temporal sites) and 300ms (on occipital sites) and showed a different pattern of task effects in each case (Itier & Neath-Tavares, 2017). Thus from the evidence reviewed so far it is not clear whether neural activity encoded within the first 200-300ms post-stimulus will show sensitivity to permit the discrimination of the basic facial expressions of emotion, and whether such sensitivity may be modified by task demands – particularly whether expressions are processed in an explicit vs implicit manner.

It is important to point out that only three prior EEG studies have investigated the effects of task on neural processing of facial expressions using within-subject designs (Itier & Neath-Tavares, 2017; Wronka & Walentowska, 2011; Rellecke et al 2012), and they have produced partially contradictory findings. Wronka & Walentowska (2011) found higher N170 for emotional than neutral stimuli but only in an explicit emotion perception task whereas Rellecke et al (2012) found higher N170 to angry faces in both explicit and implicit emotion perception tasks (they also found emotion > neutral on the later EPN component across both face tasks). Crucially none of these prior studies has investigated the *representation* of each of the basic expression categories as a function of task with MVPA.

Recently researchers have begun applying multivariate analytic techniques (MVPA) to investigate the dynamic evolution of neural representations revealed by time-sensitive neuroimaging methods, such as EEG or MEG (see Grootswagers et al 2017, for a review). Carlson et al (2013), for instance showed that visual object categories can be decoded from all-channel MEG from around 80-100ms after stimulus onset, and that onset and peak decoding occur earlier for lower (e.g. face Vs object) than higher tier categories (e.g. animate Vs inanimate). Carlson et al (2011), using all-channel MEG, further demonstrated object representations that were invariant to stimulus position by around 200ms. In addition, Cauchoix et al (2014) revealed that faces could be reliably detected within natural scenes at < 100ms and that read out was related to behavior already at 125ms post stimulus. They further argued that the decoding time-course revealed discrete stages of neural information processing. Hence, as these studies show, time-sensitive neuroimaging methods can reveal important insights into the dynamics of visual object and face processing in the brain.

In the present work we combine MVPA decoding with EEG to investigate how task shapes the neural representation of facial expression. Participants completed both an identity and an expression categorization task on the same stimulus set. One hypothesis is that visual representations of faces may be enhanced for task relevant dimensions relatively early, e.g. within the first 200ms, which would lead to enhanced processing of expression in the same expression task, and identity in the identity task at this time of processing (e.g. Schyns 1998). On the other hand, visual representations may be sensitive to the same stimulus information regardless of task at early time windows, with task sensitivity only emerging later (see e.g. Itier & Neath-Tavares, 2017; M Smith et al 2004). This view would predict that each face attribute (expression or identity) could be discriminated equally across the different tasks within the first 200ms. A third possibility (not mutually exclusive) is that facial expressions may be a specifically salient class of face information (compared to identity or gender) such that they are prioritized for neural processing even when they are not task relevant (e.g. Vuilleumier et al 2001; Anderson et al 2003). In addition to investigating how task shapes neural processing of facial expression, our analyses also examined whether sensitivity to basic facial expression categories is present on lateral occipito-temporal sites around the N170 using MVPA (see also Nemrodov et al 2016).

## Methods

### Subjects

A total of 15 right-handed (via self-report) participants (age range 18-35yrs) took part in both experiments. All participants gave written, informed consent in accordance with procedures approved by the ethics committee of the School of Psychological Sciences, Birkbeck College, University of London.

### Stimuli & Design

Participants completed both a facial expression and an identity recognition task in a single testing session lasting approximately 2 hours. We used 6 faces (3 males, 3 females) from the California Facial Expression database (Dailey et al., 2001) each posing the six basic expressions (Ekman, 1999; happy, sad, fearful, disgusted, angry, and surprised) plus neutral. The same image set of 42 images (6 identities X 7 expressions) was used for both tasks, under different categorization instructions. Images were repeated 20 times per task, for a total of 840 trials per task (120 trials per expression, 140 trials per identity), 1680 trials in total over the course of the testing session. Importantly the identity task was always completed first so as to maximize the chance of expression processing being *implicit* in this case (as our main focus here is on facial expression perception under explicit and implicit task conditions). Participants were introduced to the six new identities via pictures showing their neutral face, an associated name (Alan, Daniel, Peter, Helen, Louise, Susan) and with two fictitious facts about the person (e.g. in her spare time Louise plays guitar in a folk band).

Each trial began with a fixation cross, presented for 500ms, followed by a face stimulus from the set for 500ms followed by a 2s delay. Breaks were interposed every 140 trials in each task (for 6 short blocks per task). A longer break (minimum 10 minutes) separated the two experiments. Participants were seated at a fixed distance of 70cm from a standard CRT monitor (distance fixed by the use of a chin rest), such that the faces spanned 2.54 by 3.84 degrees of visual angle. Participants indicated their categorization choice via labeled keyboard keys. Prior to each task participants completed a short familiarization phase (42 trials) where they practiced the task and the keyboard responses.

### EEG Data Acquisition & Analysis

EEG data was recorded from 64 Ag/AgCl electrodes mounted according to the international 10:20 system in an electrode cap, electrode AFz served as ground and a single mastoid as reference. Horizontal and vertical eye movements were recorded from electrodes positioned at the outer canthi of the eyes (HEOG) and above and below the dominant eye (VEOG). Electrode impedance was lowered to <10kΩ. EEG activity was continually recorded at a sampling rate of 1000Hz, amplified between 0.1 and 300Hz. EEG data was off-line re-referenced to average reference (excluding the EOG channels), filtered between 0.01 and 40Hz and epochs created around the stimulus onset (−200ms:700ms) for each trial. The initial 200ms prior to stimulus onset acted as baseline. Trials containing artifacts were identified using standard routines in the MATLAB EEGLAB toolbox (Delorme & Makeig, 2004, 75μV threshold) and removed from further analysis (Identity task: M = 98.1 trials (SD = 97.5, 11.7%), Expression task: M=89.9 (SD = 95, 10.6%)).Channels identified as contributing excessive noise to the signal by the automatic EEGLAB routines (Kurtosis threshold 5) were removed from any further analysis (Identity task: M=4, STD=2, Expression task: M=3, STD=2).

### Multivariate Pattern Classification Analysis (MVPA)

We trained a linear classifier (Linear Support Vector Machine - SVM) to learn the mapping between a set of multivariate EEG observations of brain activity and the particular facial expression (7) or identity (6) that had been presented. We then tested the classifier on an *independent* set of test data. Importantly we decoded both expression and identity from both the expression and identity task data. We used cross-validation to assess the performance of the classifier, with a 70% train to 30% test random split of the data (see Hausfield et al., 2012; Cauchoix et al., 2014; Tsuchiya et al., 2008) repeated 20 times to form 20 cross-validation iterations (see e.g. Tsuchiya et al 2008). Importantly we sampled the same number of trials per class so as not to bias the classifier (see also Cauchoix et al 2014) and thus note if there were different numbers of trials present in each class (i.e. after EEG preprocessing) we randomly sub-sampled the same number of trials as in the smallest class from each class with a larger number of trials. The training data always consisted of single trial EEG activity patterns while the test data consisted of an average taken across the single trials comprising the test data to increase signal to noise (‘average trial analysis’, see Smith & Muckli, 2010; Petro et al., 2013). The features input to the classifier consisted of the EEG signal from a 100ms wide time window across either all electrodes OR posterior electrodes (all posterior to and including PZ, or all electrodes that were not rejected in pre-processing and excluding channels used to measure eye movements, see above). The analysis was repeated across the whole epoch of the time course by sliding the time window in 50ms offsets (thus the onset of the time windows ranged from −200 to 600ms, with 50ms offsets, giving 17 windows to cover the epoch). We note that this *shifting time-windows* method has been proposed as the best to reveal the temporal aspects of information processing in the brain (Hausfield et al 2012).

We tested whether group level decoding accuracy for each time-window was above chance by performing one-tailed t-tests against the chance level of 1/7 or 1/6 (for expression or identity, respectively; see e.g. Smith & Goodale, 2015; Walther et al. 2009; Chen et al. 2011). Control for multiple tests at multiple time windows was implemented by using the False Discovery Rate with *q* < .05. The linear SVM algorithm was implemented using the LIBSVM toolbox (Chang and Lin 2011), with default parameters (notably C=1). Note that the activity of each feature in the training data was normalized within a range of −1 to 1 prior to input to the SVM. The test data were normalized using the same parameters (min, max) as obtained from the training set normalization in order to optimize the classification performance (see Chang and Lin 2011; Smith & Muckli, 2010; Smith & Goodale, 2015; Vetter et al., 2014).

### Brain Behavior Correlations

From the decoding analysis we also extracted a confusion matrix (see e.g. Greening et al 2018; Smith & Goodale 2015; Vetter et al 2014) at each time-window that reveals the pattern of errors made by the classifier (i.e. it reveals the probability of a correct response per class plus the probability of incorrectly assigning each other class, for all classes). For each participant in turn, we correlated the errors in decoding (i.e. the non-diagonal elements of the confusion matrix) for each time-window with the errors that participant made in each categorization task. We used Spearman correlation to mitigate against the risk of outliers affecting the Pearson correlation coefficient (see e.g. Pernet et al., 2012; Greening et al 2018). We Fisher transformed the Spearman rho values to allow averaging across participants.

### Spatially Resolved Classification Analysis

In order to address which electrodes drove classification performance at what time-windows, we performed an additional classification analysis (see e.g. Nemrodov et al., 2016). In this analysis, we ran the classification analysis as described above except that the analysis was performed independently for each electrode in turn. Thus the features input to the classifier were the EEG signal amplitudes over a given time-window for a given electrode (see Nemrodov et al., 2016). This allowed us to create a scalp map which shows a spatial map of where facial expression information can be read out at specific time-windows. We ran this analysis using the same time-windows as the main analyses (i.e. 100ms length, 50ms offset). We chose this method, as opposed to visualizing the weights of the multi-channel classifier, because the weights of any linear classifier that takes into account interactions between voxels, such as a linear SVM, are typically hard to interpret when projected back into brain space (see Pereira et al 2009; but see also the correction method proposed in Haufe et al., 2014).

### Low Level Image Analysis

We computed pixel-wise Euclidean distances across our images to reveal which face property may be more easily decoded via the low level image statistics (see also Greening et al., 2018). In particular, for each expression category in turn, we computed the difference across each pair of the 6 identities used (N=15 pairs). This reveals how well face identity can be discriminated from the image pixel values. We then computed, for each identity in turn, the difference across each pair of expressions (N=21 pairs). This reveals how well facial expression can be discriminated from the image pixel values. This analysis is useful because it is simple and does not rely on a complicated algorithm to determine discriminability.

## Results

### Analysis of Behavior

Average performance on the expression task reached 69% correct (SD = 10) which was subjectively much lower than on the identity task, where performance was higher at 92% correct (SD = 9). As the chance levels were different we did not explicitly compare performance across tasks. The confusion matrices underlying this performance can be seen in Figure 1. For the expression categorization task, a one-way repeated measures ANOVA revealed that the effect of emotion was highly significant, *F*(6,84) = 13.72, *p* < .001, = 0.495. Follow-up paired sample t-tests with a Bonferonni correction revealed that happy faces were recognized significantly more accurately than fearful, disgusted, angry and sad faces (all *t*’s > 4.31, *all p*’s < .0007, all *d*’s >= 1.11) while there was a trend for happy faces to be better recognized than both surprised (*p* = .0043, *d* = .88) and neutral faces (*p* = .0055, *d* = .85), which did not survive multiple comparison correction. Neutral faces were recognized significantly better than fearful faces (*t*(14) = 5.23, p = .0001, *d* = 1.34) and displayed a trend for better recognition than sad (*p* = .0097, *d* = .77) and disgusted faces (*p* = .042, *d* = .58). Disgusted, surprised, angry and sad faces were all better recognized than fearful faces (all *t*’s > 3.90, all *p*’s < .0016, all *d*’s >= 1.01). Thus agreeing with previous research, happy faces were generally the best recognized facial expression (Smith & Schyns, 2009; Smith & Rossit, 2018).

**Figure 1:**
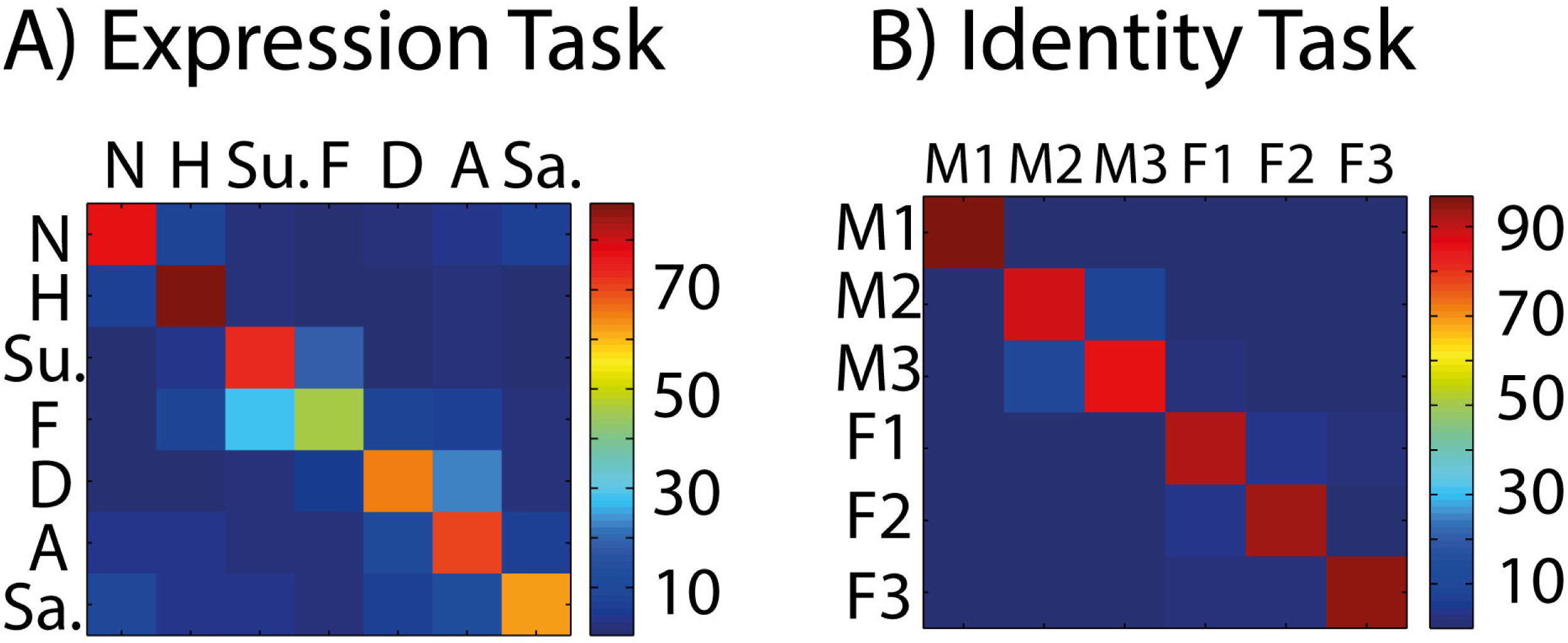
Human Categorization Performance. A. **Confusion matrix pertaining to Expression Task. Rows represent expression category and columns represent the response chosen by the participant. The diagonal hence represents correct responses, and the off diagonal errors. The colour scale indicates the percentage of times a particular stimulus and response were chosen.**
B. **As in A but for the Identity Task. Rows and columns here hence represent identity (1-6).**

### Decoding Analyses

We computed decoding performance across a sliding time window (100ms long, 50ms offset: see Methods) to reveal how well facial expression (or identity) category can be read out from multi-channel EEG activity, under both explicit and implicit processing conditions (Figure 2). We computed performance both across all electrodes and for a posterior subset of electrodes (see Methods) as a proxy for those more related to visual processing. We focus on the results obtained with the posterior subset below, but note a similar pattern emerged using all electrodes (see Supplementary Material).

**Figure 2:**
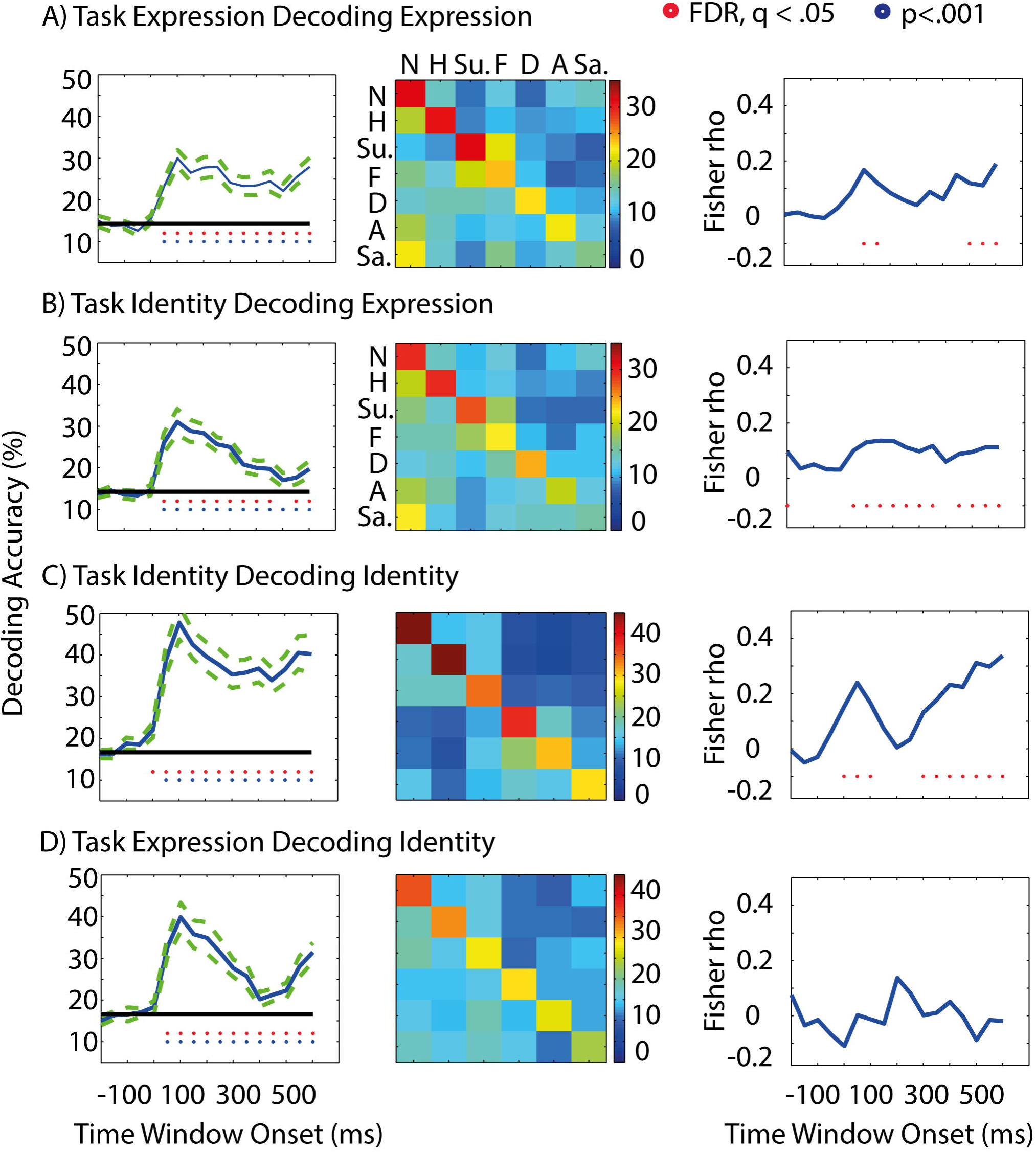
Decoding of Facial Expression and Facial Identity in Explicit and Implicit Tasks. A. **Left Panel: Decoding of facial expression in explicit task (while participants categorized the faces by expression) for each time-window (100ms wide, 50ms offsets, covering the whole epoch). Red stars represent significant decoding FDR corrected q < .05 (chance ~14%, solid black line). Blue circles represent uncorrected p < .001. Middle panel: Confusion matrix pertaining to the decoding results at left. Rows represent expression category and columns the response chosen by the decoder. The diagonal hence represents correct decisions and the off-diagonal reflects the errors made by the decoder. Right Panel: Correlation between errors in human behavioral categorization of expression and errors in decoding.**
B. **As above but for implicit decoding of facial expression (i.e. decoding identity while participants categorized the faces by identity).**
C. **As above but for explicit decoding of identity (i.e. decoding identity while participants categorized the faces by identity). Chance 16.67%.**
D. **As Above but for implicit decoding of identity (i.e. decoding identity while participants categorized the faces due to expression). Chance ^~^16.67%.**.

### Explicit Decoding of Expression

For decoding of facial expression under explicit conditions (i.e. when participants perform an expression categorization task), decoding was first significant within the 50-150ms time window and extended across the whole epoch (Figure 2A). We note, moreover, that decoding peaked both in an earlier (100-200ms) and later time window (600-700ms) with a trough in between. Thus facial expression information contained within the EEG signal does not simply increase monotonically with time, which highlights the possibility of several discrete processing stages being revealed in the decoding time-course (see also Cauchoix et al 2014). Figure 2A, second panel, depicts the confusion matrix underlying this classification performance (averaged across all time windows that led to FDR q <.05) and shows that most expressions could be well discriminated except sadness.

### Implicit Decoding of Expression

We repeated the same analyses (i.e. decoding expression) from the data acquired while participants performed the identity task (Figure 2B). Decoding was again first significant within the 50-150ms time window and peaked at 100-200ms. Decoding remained significant until 450-550ms (and was again significant at 600-700ms). There was, however, no pronounced later decoding peak in this analysis, unlike in the explicit case above, and a steep decline after the initial peak before a small rebound at the end of the epoch. Thus the maintenance of face information seen in explicit decoding above may reflect the need to keep the information available for later read-out. Figure 2B, second column, depicts the relevant confusion matrix highlighting that again most expressions are well discriminated, except sadness.

### Explicit Decoding of Identity

Reliable decoding of identity under explicit task conditions began within the 0-100ms time window and remained significant across the whole epoch (Figure 2C). As in explicit expression decoding, the time course of decoding contained two separate peaks, one in an earlier (100-200ms) and one in a later (550-650ms) time window, with a trough in between. Figure 2C, second column, depicts the confusion matrix underlying this classification, and shows that all identities (6) could be well discriminated. Interestingly the gender of the stimuli is reflected in the confusability of the male identities (1-3) with one another but not with the female identities (4-6), which also tend to be confused with one another.

### Implicit Decoding of Identity

Reliable decoding of identity began slightly later within the 50-150ms time window, again peaked at 100-200ms time-window, and extended across the entire epoch (Figure 2D). However, the decoding magnitude dropped sharply after peaking (from ~40 to ~20%), but identity read-out rebounded towards the end of the epoch (600-700ms, ~30%). Figure 2D, second column, shows that while most identities are relatively well discriminated, the confusability due to the gender of the stimuli is absent when identity is perceived in an implicit manner. Thus the gender confusability found during explicit identity decoding is likely a high level effect due to the specific task demands of categorizing identity.

### Analysis of Peak Decoding

We compared peak decoding magnitude (computed per participant across time-window onsets of 50, 100 and 150ms; see Methods) across explicit Vs implicit processing for expression and identity tasks independently (as each task has a different chance level). These analyses revealed no change in peak decoding of expression as a function of task (*t*(14) = 1.18, *p* = .26, two-tailed; *d* = 0.30; Explicit = 32%, Implicit = 34%) but a significant decline in decoding of identity in the implicit task (*t* (14) = 3.96, *p* = .0014, two-tailed; *d* = 1.02; Explicit = 50%, Implicit = 41%). Thus these results reveal that identity information is not as well represented during implicit conditions whereas expression information is equally well represented independent of the particular processing condition (i.e. explicit Vs implicit). We note that the same pattern of results is present if we use all electrodes rather than just the posterior subset – see Supplementary Figure 1.

### Brain Behavior Correlations

We next computed, per participant, the Spearman correlation between errors in the brain decoding (off diagonal elements of confusion matrix, see Figure 2, second column) at each time-window, with the errors participants made in the relevant categorization task. Figure 2 (Third Column) shows the resulting correlations averaged across participants. These analyses thus depict when behaviorally relevant information is present in the decoding observed (see e.g. Walther et al., 2009). Note we computed these correlations for each face attribute (expression or identity) under both explicit and implicit processing conditions. Our rationale for computing correlations even under implicit processing was that this would give a measure of the quality of higher level information that is available under these conditions. There were significant correlations for both expression and identity decoding under explicit processing conditions, in both earlier (Expression onset: 100-200ms; Identity onset: 50-150ms) and later (Expression onset: 500-600ms; Identity onset: 300-400ms) time-windows, resembling the two peaked nature of the decoding time course. However, under implicit processing conditions, correlations only reached significance for decoding expression but not for identity, and were relatively sustained across most of the epoch. Thus, behaviorally relevant information is present for expression regardless of the specific processing conditions but not for identity. Again, similar results were present using all available electrodes (see Supplementary Figure 1).

### Analysis of Low Level Features

We computed pixel-wise Euclidean distances across identities (within an expression) and across expressions (within identities) categories to determine which would be more easily discriminable in principle (i.e. have greater distance on average; see Methods). These analyses revealed that identity information is more readily discriminable from the low level visual input than is expression information (i.e. has larger Euclidean distance; Mean Identity 7.17 × 10^3^, Mean Expression = 5.21 × 10^3^, see Supplementary Table 1). This was true both on average, and for each expression and identity sub-category (i.e. 7 expressions and 6 identities: See Supplementary Table 1). Thus finding significant correlations on the implicit task for decoding of expression but not identity is intriguing as if these correlations merely reflected low level feature processing, one would have expected to find robust correlations present also for identity on the implicit task. In addition, the fact that identity is more easily discriminable from the image pixel values likely contributes to the overall higher accuracy for decoding identity versus expression from our face set.

### Spatially-Resolved Decoding of Face Information

In order to determine which specific electrodes may be contributing to the successful decoding reported above, we re-ran our classification analyses independently per electrode. Figures 3 and 4 shows the results of these analyses. Across both tasks, and both face attributes, information first became reliably decodable at 50-150ms post stimulus, matching well the onset of decoding for the multi-channel analyses reported earlier.

**Figure 3:**
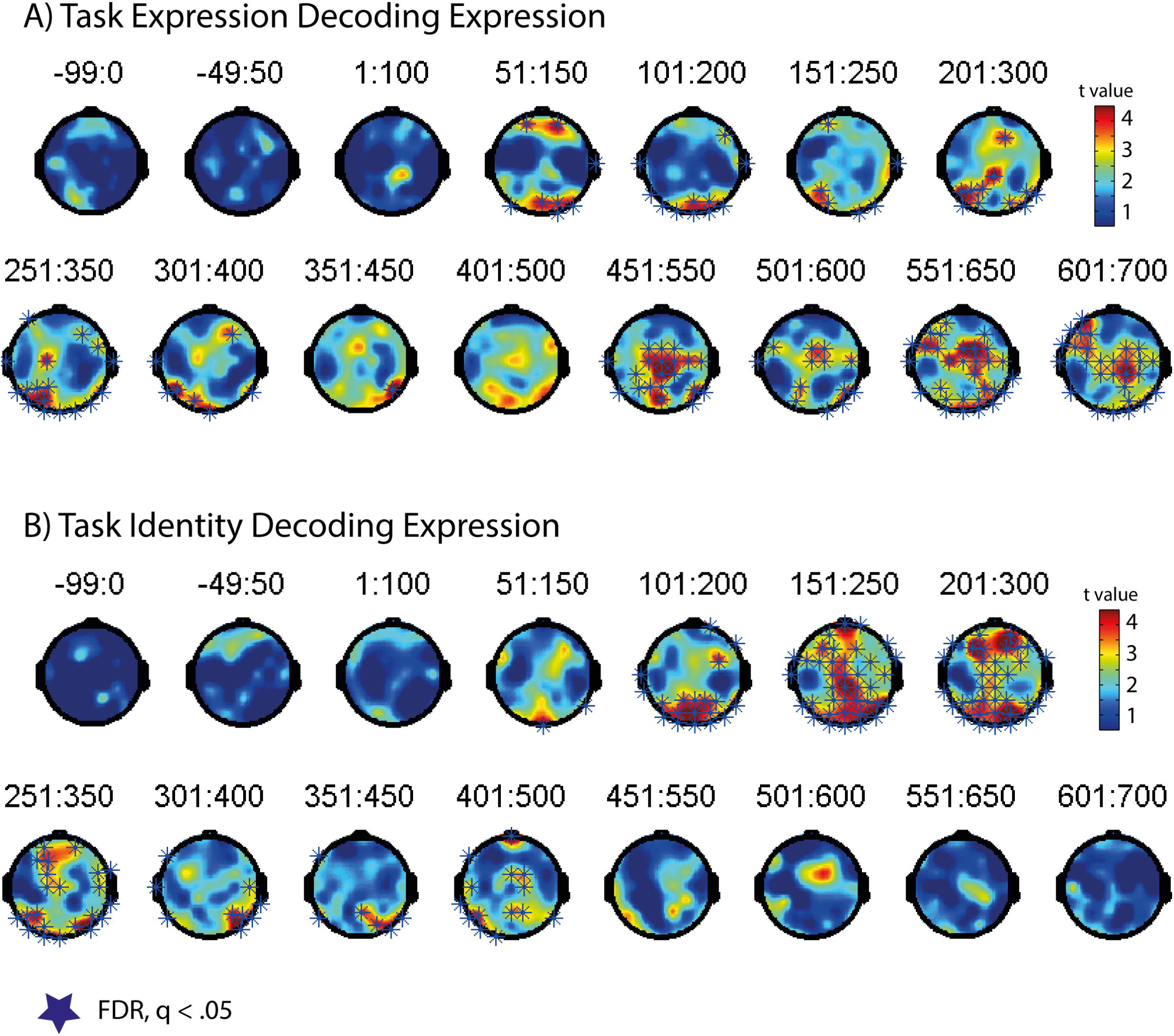
Spatially-Resolved Decoding of Facial Expression in Explicit and Implicit Tasks. A. **Decoding of facial expression in the explicit task. Each map shows the t-values for whether decoding is above chance at each electrode for a given time-window. Blue stars indicate FDR corrected q < .05.**
B. **As in A, but for decoding of facial expression in the implicit (identity) task.**

**Figure 4:**
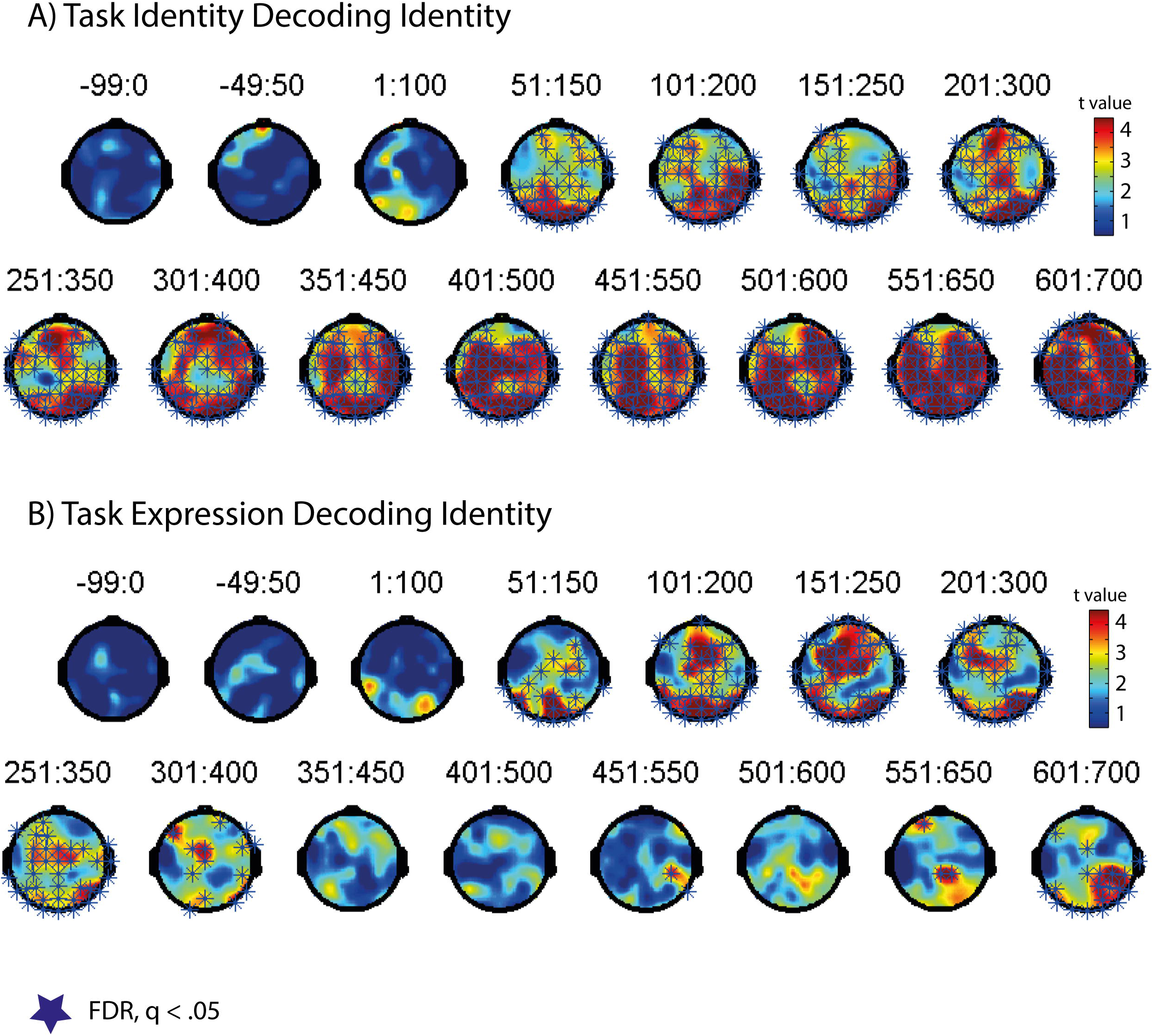
Spatially-Resolved Decoding of Facial Identity in Explicit and Implicit Tasks. A. **Decoding of facial identity in the explicit (identity) task. Each map shows the t-values for whether decoding is above chance at each electrode for a given time-window. Blue stars indicate FDR corrected q < .05.**
B. **As in A, but for decoding of facial identity in the implicit (expression) task.**

In explicit decoding of expression (Figure 3A), posterior and frontal electrodes showed significant decoding initially. Central sites showed significant decoding around the time-window of the P3 component (200-300ms time-window). There was then very sparse (350-450ms) or no decoding (400-500ms time-window) before decoding rebounded strongly from 450-550ms until the end of the epoch. At this later stage, information was maximally present at central, frontal and to a lesser extent posterior electrodes, presumably reflecting response related activity.

In implicit decoding of expression (Figure 3B), initially (51-150ms, 101-200ms) posterior electrodes showed the most robust decoding. Between 150-300ms robust decoding was present at both central and frontal sites, in addition to posterior electrodes. Sparse and weaker decoding was present across 301-400ms (mainly right posterior and some central and frontal sites) before an absence of decoding from 450-550ms onwards.

For explicit decoding of identity (Figure 4A), initially posterior sites showed most robust decoding (51-150ms) although some frontal sites already showed significant decoding as well. This profile changed (201-300ms) to include central sites more strongly in addition to frontal and posterior. From 351-450 time window onwards, very robust decoding was present throughout most of the scalp. For implicit decoding of identity (Figure 4B), on the other hand, decoding was robustly present at posterior sites initially (50-150ms) and then became more robust at frontal and central sites (101-200 and 151-250ms time-windows). Decoding was largely absent from 350-450ms onwards but was significant again at 551-650ms onwards.

Hence decoding of either identity or expression in the implicit task followed the same pattern: initially most robust decoding at posterior sites which then moves to central and frontal sites. Explicit decoding, on the other hand, displayed reliable decoding at posterior and frontal sites already in the first time-window (50-150ms). In addition, these analyses match well with the multi-channel analyses reported earlier in terms of temporal profile of available face information.

### Is information about facial expression present at occipito-temporal sites around the N170 peak

In order to address this question, we looked at which lateral occipito-temporal electrodes revealed above chance decoding in the spatially resolved decoding analyses. From looking at Figures 3 and 4, reliable decoding of identity was generally present in the majority of such electrodes across both tasks around the N170 (time-windows of 100-200 and 150-200ms). Decoding of expression was also present in a subset of these same electrodes (see Figure 2A & 2B 100-200ms time-window), with more such electrodes being significant in *implicit* than in *explicit* decoding. Thus our data clearly show that lateral OT sites do contain information about facial expressions around the time-window of the N170, in addition to information about identity.

## Discussion

In the present study we investigated how task shapes representations of facial expression and facial identity, using EEG with MVPA. Our findings show that peak decoding for expression is not affected by task set whereas peak decoding of identity is affected by task set. Further our results demonstrate distinct decoding time-courses for explicit Vs implicit processing of face categories, each displaying evidence of discrete processing stages. We report reliable correlations of the errors in neural decoding with errors in human categorization at both earlier and later periods during explicit tasks but only for expression in an implicit task at earlier time points. Spatially-resolved decoding further revealed distinct patterns of informative electrodes in explicit Vs implicit tasks. Finally we provide novel evidence that lateral occipito-temporal sites encode information about both facial identity *and* facial expression around the time-window of the N170.

### Implicit task context weakens identity but not expression decoding

We report reliably greater decoding of identity when identity is task relevant, within the first 300ms (the time-window including the P1 and the N170 visual responses). However, for expression, we did not find any difference as a function of task. These findings are consistent with two accounts of how task shapes visual processing of face categories. First, the change of performance for identity may be due to top down attentional mechanisms boosting processing of identity when it is a task relevant dimension comparable to effects reported in early visual areas for simpler dimensions (see e.g. Maunsell, 2015) and to those recently reported in face areas for identity (Gratton et al., 2013; see also Kay & Yeatman, 2017). This is also consistent with the greater BOLD responses observed in ventral regions for attending to identity Vs expression, whereas STS shows the opposite pattern (see Hoffman & Haxby 2000). Such effects of attention on high level visual regions may be driven by regions in the IPS (Kay & Yeatman, 2017).

If we assume the same account applies to expression perception, then this would imply that the failure to find a significant difference for expression across tasks, is due to the absence of such top down mechanisms operating during explicit expression decoding within the first 300ms. That is, expression perception would be determined by the same bottom-up stimulus processing in both cases. However there are several reasons why we do not think this is the case: First, top-down influences on emotion perception are known to occur and generate feedback to visual cortex (e.g. Furl et al 2013; Vuilleumier et al 2004). Second, we found reliable correlations of the implicit neural representation of emotion with participants’ explicit emotion perception–implying that a relatively rich representation of emotion is constructed even when emotion is non-task relevant. In fact, a long-standing body of work on emotion and attention, has revealed that it is difficult to ignore the emotion presented in sensory stimuli, unless attention is very highly loaded on a different attribute: something that would be rather unlikely given the present experimental design (see e.g. Pessoa & Ungerleider, 2004; Phelps et al 2006).

Hence we argue that due to the evolutionary importance of facial expressions as high-value signals - that transmit information about the mental states, intentions and environment of the expresser (Darwin 1872; Fridlund, 1994; Matsumoto, 2001) - relative to that of facial identity signals – our results may actually reveal no difference between expression decoding in explicit and implicit task contexts *because* emotions are preferentially processed even in implicit task contexts, whereas facial identities are not (perhaps particularly in the case of very simply and recently learned identities as used in the present experiment). While previous neuroimaging studies have revealed that different brain areas may be active in implicit Vs explicit emotion perception, overall the pattern is still unclear and in need of further investigation (e.g. Critchley et al., 2000; Chi-Hua Chen et al., 2006; Hariri et al 1999; Lange et al., 2006; but see also Gur et al 2002).

A recent fMRI study (Dobs et al 2018), however, directly addressed how task shapes neural representation of facial expression and identity revealed that surprisingly, both early visual areas (V1-V4) and STS discriminated facial expressions better when participants performed an expression task, while both FFA and STS showed better decoding of identity in an identity task. Thus in this case, clear effects of task shaping visual representations were found for both face attributes, albeit in different brain areas. While the Dobs et al study is admirable for equating task difficulty and stimulus differences precisely across key comparisons, the study used quite a limited set of facial expressions (just angry and happy) and facial identities (two females), and employed stimuli that are clearly artificial. Hence it is unclear to what extent the findings are truly generalizable across different expression categories in the different task contexts (here, we use the full set of six basic expressions plus neutral, and six identities comprising both males and females). Future studies, in any case, will be necessary to reveal the extent to which facial expressions may be preferentially encoded independent of task.

### Identity and Expression were maximally decodable within a 100-200ms time-window over posterior electrodes

Despite the key difference noted above in how task set affected peak decoding for expression vs identity, there were notable broad similarities in the decoding time-courses for each face attribute when perceived in explicit Vs implicit task contexts. First, decoding was maximal in the 100-200ms time-window post-stimulus irrespective of task and face attribute. Thus for the posterior set of electrodes used in our main analyses (posterior to and including Pz), this demonstrates that maximal face information about identity and expression is present in the 100-200ms post-stimulus time-window. Previous work has revealed that read-out of face exemplar information from whole brain MEG is present across an ~ 80-200ms time-period (Carlson et al 2013; see also Carlson et al., 2011). In addition, Nemrodov et al (2016), revealed a clear peak in decoding around 150ms for decoding both face identity and gender from all scalp EEG (initial smaller peak at 70ms). Thus the present findings are in broad agreement with the limited previous literature that has attempted to decode fine-grained properties about human faces from high temporal resolution neuroimaging techniques. Importantly the present study goes beyond this literature by systematically quantifying the information available about both the expression and identity of human faces across time, in both explicit and implicit task contexts.

### Explicit and Implicit Tasks produce different decoding time-courses and spatial patterns of informative electrodes

Decoding face attributes under explicit task conditions led to sustained decoding across the whole decoding time-course, beginning within the 0-100 or 50-150ms time-windows, and then declining somewhat before plateauing (~200-500ms) before rebounding at the end of the epoch (550-700ms). On the other hand, decoding in implicit task contexts led to more focal burst of decoding within an early time-period (peaking 100-200ms) but with a steep decline of decoding through an intermediate period (~150-500ms) before rebounding at the end of the epoch (~550-700ms). In agreement with previous work (Cauchoix et al., 2012), this reveals that the quality of face information changes across the epoch and does not simply increase monotonically as a function of time. In particular, under explicit conditions, the first phase is characterized by accumulation of face information (0-200ms) which may reflect low level feature processing in early visual areas. The second phase (100 – 450ms) shows a plateau in both tasks which may indicate maintenance of the available face information in higher level brain areas important in face processing such as OFA, FFA and STS. The third phase (450-700ms) displays a rebound which may indicate the involvement of feedback from frontal or motor planning regions to visual cortex for response read out. This fits with the spatially resolved decoding analyses (Figures 3 and 4) revealing robust decoding at central and frontal sites, in addition to posterior sites, across this time period.

For implicit processing, on the other hand, the first phase again revealed the same rapid accumulation of face information over the first few hundred ms. However here the second phase (~150 – 500ms) did not reflect a plateau but rather revealed a steep decrease in the face information available across time in both tasks. This may reflect the fact that implicit processing does not require that the face information be maintained in higher-level face areas in order for explicit categorization. This is also shown by the less robust decoding in posterior electrodes around 350-600ms in the spatially resolved decoding (See Figures 3B & 4B). Intriguingly the availability of face information again rebounded at the end of the epoch (~550-700ms). The fact that decoding rebounds across both modes of face processing, suggests that information about both categories is made more accessible around the time of response read-out, and thus that the availability of information at this time does not just solely reflect response requirements of the task.

### The N170 carries information about both facial identity and facial expression

It has been unclear from previous work whether the N170 component is sensitive to facial expression, in addition to identity. Some studies have found effects of emotion on the N170 (Batty & Taylor, 2003; Leppanen et al., 2007, 2008; Schyns et al., 2007) whereas others have not found such effects (Eimer & Holmes, 2002; Eimer & Holmes, 2007; Pourtois et al 2005; Neath-Tavares & Itier, 2016)). Here we demonstrate that lateral OT sites around the time of N170 do carry information about facial expression categories when expression is perceived both in an explicit and implicit task context. While decoding of identity was arguably more robustly present across all analyses at such sites, decoding of expression was clearly present. As multivariate techniques, such as the linear decoding that we use here, are known to be more sensitive or powerful than univariate techniques, it is likely that information about facial expression is generally present in the EEG signal around the N170, but that it is more weakly encoded than is information about identity. Future experiments, where behavioral performance is equalized would be necessary to formally test this hypothesis. In addition, possible future claims for stronger read out of identity face information than expression would also need to take into account the fact that identity is typically much more easily decoded from the low level image statistics than is facial expression.

## Conclusion

In summary, the present study reveal differences and commonalities in the processing of face categories under explicit Vs implicit task contexts and suggest that facial expressions are processed to a richer degree even under implicit processing conditions, consistent with prior work indicating the relative automaticity by which emotion is processed. Our results further demonstrate evidence of discrete processing stages in the dynamically evolving representation of two important face categories in visual cortex together with distinct patterns of informative electrode sites across explicit and implicit modes of face perception. Finally our work shows the utility in applying multivariate decoding analyses to EEG for revealing the dynamics of face perception.

## References

Anderson AK, Christoff K, Panitz D, De Rosa E, Gabrieli JDE. (2003). Neural correlates of the automatic processing of facial threat signals. Journal of Neuroscience, 23, 5267–5633.

Batty, M., & Taylor, M. J. (2003). Early processing of the six basic facial emotional expressions. Cognitive Brain Research, 17(3), 613–620. doi:http://dx.doi.org/10.1016/S0926-6410(03)00174-5

Carlson TA, Hogendoorn H, Kanai R, Mesik J, Turret J. (2011). High temporal resolution decoding of object position and category. Journal of Vision, 11, 9.

Carlson TA, Tovar DA, Alink A & Kriegeskorte N. (2013). Representational dynamics of object vision: The first 1000ms. Journal of Vision, 13, 1.

Cauchoix, M., Barragan-Jason, G., Serre, T., & Barbeau, E. J. (2014). The neural dynamics of face detection in the wild revealed by MVPA. J Neurosci, 34(3), 846–854. doi:10.1523/JNEUROSCI.3030-13.2014

Chang CC, Lin CJ. 2011. LIBSVM: a library for support vector machines. ACM Transactions on Intelligent Systems and Technology 2:27.

Chen Y, Namburi P, Elliott LT, Heinzle J, Soon CS, Chee MW, Haynes JD. 2011. Cortical surface-based searchlight decoding. Neuroimage. 56:582–592.

Critchley, H., Daly, E., Phillips, M., Brammer, M., Bullmore, E., Williams, S., … Murphy, D. (2000). Explicit and implicit neural mechanisms for processing of social information from facial expressions: A functional magnetic resonance imaging study. Hum Brain Mapp, 9(2), 93–105. doi:10.1002/(SICI)1097-0193(200002)9:2<93::AID-HBM4>3,0.CO;2-Z

Darwin C. (1872). The expression of emotion in man and animals. New York, NY: Oxford University Press.

Delorme, A. & Makeig, S. (2004). EEGLAB: an open source toolbox for analysis of single-trial EEG dynamics including independent component analysis. Journal of Neuroscience Methods, 134(1), 9–21.

Desimone R, Duncan J. (1995). Neural mechanisms of selective visual attention. Annual Reviews Neuroscience, 18, 193–222.

Dobs K, Schultz J, Bulthoff I, Gardner JL. (2018). Task-dependent enhancement of facial expression and identity representations in human cortex. Neuroimage, 172, 689–702.

Eimer, M., & Holmes, A. (2002). An ERP study on the time course of emotional face processing. NeuroReport, 13(4), 427–431.

Eimer, M., & Holmes, A. (2007). Event-related brain potential correlates of emotional face processing. Neuropsychologia, 45(1), 15–31. doi:10.1016/j.neuropsychologia.2006.04.022

Eimer M. (2011). The face-sensitive N170 component of the event-related brain potential. In Calder AJ et al (eds). The Oxford Handbook of Face Perception. Oxford University Press.

Engell AD, Haxby JV (2007) Facial expression and gaze-direction in human superior temporal sulcus. Neuropsychologia 45:3234–3241.

Fridlund A. (1994). Human Facial Expression: An evolutionary view. San Diego, CA: Academic Press.

Fusar-Poli P, Placentino A, Carletti F, Landi P, Allen P, Surguladze S, Benedetti F, Abbamonte M, Gasparotti R, Barale F, Perez J, McGuire P, Politi P (2009) Functional atlas of emotional faces processing: a voxel-based meta-analysis of 105 functional magnetic resonance imaging studies. J Psychiatry Neurosci 34:418–432.

Furl N, Henson RN, Friston KJ, Calder AJ (2013) Top-down control of visual responses to fear by the amygdala. J Neurosci 33:17435–17443.

Gilbert CD, Li W. (2013). Top-down influences on visual processing. Nature Reviews Neuroscience, 14, 350–363.

Gratton C, Sreenivasan KK, Silver MA, D-Epsposito M. (2013). Attention selectively modifies the representation of individual faces in the human brain. Journal of Neuroscience, 33, 6979–6989.

Greening, S. G., Mitchell, D. G. V., & Smith, F. W. (2018). Spatially generalizable representations of facial expressions: Decoding across partial face samples. Cortex, 101, 31–43. doi:https://doi.org/10.1016/j.cortex.2017.11.016

Grootswagers, T., Wardle, S. G., & Carlson, T. A. (2017). Decoding Dynamic Brain Patterns from Evoked Responses: A Tutorial on Multivariate Pattern Analysis Applied to Time Series Neuroimaging Data. Journal of Cognitive Neuroscience, 29(4), 677–697. doi:10.1162/jocn_a_01068

Gur RC, Schroeder L, Turner T, McGrath C, Chan RM et al. (2002). Brain activation during facial emotion processing. Neuroimage, 16, 651–662.

Hariri, A. R., Bookheimer, S. Y., & Mazziotta, J. C. (2000). Modulating emotional responses: effects of a neocortical network on the limbic system. NeuroReport, 11(1), 43–48.

Hausfield L, De Martino F, Bonte M & Formisano E. (2012). Pattern analysis of EEG responses to speech and voice: influence of feature grouping. Neuroimage, 59, 3641–3651.

Haxby JV, Hoffman EA, Gobbini MI (2000) The distributed human neural system for face perception. Trends Cogn Sci 4:223–233.

Haufe S, Meinecke F, Gorgen K, Dahne S, Haynes JD et al. (2014). On the interpretation of weight vectors of linear models in multivariate neuroimaging. Neuroimage, 87, 96–110.

Hinojosa, J. A., Mercado, F., & Carretié, L. (2015). N170 sensitivity to facial expression: A meta-analysis. Neuroscience & Biobehavioral Reviews, 55, 498–509. doi:http://dx.doi.org/10.1016/j.neubiorev.2015.06.002

Hoffman EA, Haxby JV. (2000). Distinct representations of eye gaze and identity in the distributed human neural system for face perception. Nature Neuroscience, 3, 80–84.

Kay KN, Yeatman, JD. (2017). Bottom-up and top-down computations in word- and face-selective cortex. eLife, 6:e22341 doi: 10.7554/eLife.22341.

Leppänen, J. M., Hietanen, J. K., & Koskinen, K. (2008). Differential early ERPs to fearful versus neutral facial expressions: A response to the salience of the eyes? Biological Psychology, 78(2), 150–158. doi:https://doi.org/10.1016/j.biopsycho.2008.02.002

Li W, Piech V, Gilbert CD. (2004). Perceptual learning and top-down influences in primary visual cortex. Nature Neuroscience, 7, 651–657.

Luck SJ, Woodman GF, Vogel EK. (2000). Event-related potential studies of attention. Trends in Cognitive Sciences. 4(11), 432–440.

Luo W, Feng W, He W, Wang N, Luo Y. (2010). Three stages of facial expression processing: ERP study with rapid serial visual presentation. Neuroimage, 49, 1857–1867.

Maunsell, JHR. (2015). Neuronal mechanisms of visual attention. Annual Review Vision Science, 1, 373–391.

Matsumoto D, Hwang RS. (2011). Judgments of facial expressions of emotion in profile. Emotion, 11, 1223–1229.

Neath-Tavares, K. N., & Itier, R. J. (2016). Neural processing of fearful and happy facial expressions during emotion-relevant and emotion-irrelevant tasks: A fixation-to-feature approach. Biological Psychology, 119(Supplement C), 122–140. doi:https://doi.org/10.1016/j.biopsycho.2016.07.013

Nemrodov, D., Niemeier, M., Mok, J. N. Y., & Nestor, A. (2016). The time course of individual face recognition: A pattern analysis of ERP signals. Neuroimage, 132, 469–476. doi:https://doi.org/10.1016/j.neuroimage.2016.03.006

Peelen MV, Fei Fei L, Kastner S. (2009). Neural mechanisms of rapid natural scene categorization in human visual cortex. Nature, 460, 94–97.

Pereira F, Mitchell T, Botvinick M. (2009). Machine learning classifiers and fMRI: a tutorial overview. Neuroimage, 45, S199–S209.

Pernet CR, Wilcox R, Rousselet GA (2012) Robust correlation analyses: false positive and power validation using a new open source Matlab toolbox. Frontiers in psychology 3.

Petro LS, Smith FW, Schyns PG, Muckli L (2013) Decoding face categories in diagnostic subregions of primary visual cortex. Eur J Neurosci 37:1130–1139.

Pessoa L, Ungerleider LG. (2004). Neuroimaging studies of attention and the processing of emotion-laden stimuli. Prog. Brain Research, 144, 171–182.

Phelps EA, Ling S, Carrasco M. (2006). Emotion facilitates and potentiates the perceptual benefits of attention. Psychological Science, 17, 292–299.

Pourtois G, Dan ES, Grandjean D, Sander D, Vuilleumier P. (2005). Enhanced extrastriate visual response to bandpass: time course and topographic evoked potentials mapping. Human Brain Mapping 26, 65–79.

Recio G, Wilhelm O, Sommer W, Hildebrandt A. (2017). Are event-related potentials to dynamic facial expressions of emotion related to individual differences in the accuracy of processing facial expressions and identity? Cogn. Affect. Behav. Neurosci. 17, 364–380. 10.3758/s13415-016-0484-6

Rellecke J, Sommer W, Schacht A. (2012). Does processing of emotional facial expressions depend on intention? Time-resolved evidence from event-related brain potentials. Biological Psychology, 90, 23–32.

Rossion B, Jacques C. (2012). The N170: Understanding the time course of face perception in the human brain. In: Luck, S.J., Kappenman, E.S. (Eds.), The Oxford Handbook of Event-Related Potential Components. Oxford University Press, Oxford, pp. 115–141.

Schyns PG. (1998). Diagnostic recognition: task constraints, object information, and their interactions. Cognition, 67, 147–179.

Schyns, P. G., Petro, L. S., & Smith, M. L. (2007). Dynamics of Visual Information Integration in the Brain for Categorizing Facial Expressions. Current Biology, 17(18), 1580–1585. doi:10.1016/j.cub.2007.08.048

Smith E, Weinberg A, Moran T, Hajcak G. (2013). Electrocortical responses to NIMSTIM facial expressions of emotion. Int. J. Psychophysiology, 88, 17–25.

Smith FW, Schyns PG. (2009). Smile through your fear and sadness: transmitting and identifying facial expression signals over a range of viewing distances. Psychological Science, 20(10), 1202–1208.

Smith FW, Muckli L (2010) Nonstimulated early visual areas carry information about surrounding context. Proc Natl Acad Sci U S A 107:20099–20103

Smith FW, Goodale MA (2015) Decoding visual object categories in early somatosensory cortex. Cereb Cortex 25:1020–1031.

Smith FW, Rossit S. (2018). Identifying and detecting facial expressions of emotion in peripheral vision. PLOS ONE, 13: e0197160, doi: 10.1371/journal.pone.0197160

Smith, M. L., Gosselin, F., & Schyns, P. G. (2004). Receptive Fields for Flexible Face Categorizations. Psychological Science, 15(11), 753–761. doi:10.1111/j.0956-7976.2004.00752.x

Smith, M. L. (2012). Rapid Processing of Emotional Expressions without Conscious Awareness. Cerebral Cortex, 8, 1748–1760.

Turano, M. T., Lao, J., Richoz, A.-R., Lissa, P. d., Degosciu, S. B. A., Viggiano, M. P., & Caldara, R. (2017). Fear boosts the early neural coding of faces. Social Cognitive and Affective Neuroscience, 12(12), 1959–1971. doi:10.1093/scan/nsx110

Tsuchiya N, Kawasaki H, Oya H, Howard MA, 3rd, Adolphs R (2008) Decoding face information in time, frequency and space from direct intracranial recordings of the human brain. PLoS One 3:e3892.

Vetter, P., Smith, F.W., & Muckli, L. (2014). Decoding sound and imagery content in early visual cortex. Current Biology. 24, 1256–1262.

Vuilleumier P, Armony J, Driver J, Dolan RJ. (2001). Effects of attention and emotion on face processing in the human brain: an fMRI study. Neuron, 30, 829–841.

Vuilleumier P, Richardson MP, Armony JL, Driver J, Dolan RJ (2004) Distant influences of amygdala lesion on visual cortical activation during emotional face processing. Nat Neurosci 7:1271–1278.

Vuilleumier P, Pourtois G. (2007). Distributed and interactive brain mechanisms during emotion face perception: Evidence from functional neuroimaging. Neuropsychologia 45, 174–194.

Walther DB, Caddigan E, Fei-Fei L, Beck DM. 2009. Natural scene categories revealed in distributed patterns of activity in the human brain. J. Neurosci. 29:10573–10581.

Wegrzyn M, Riehle M, Labudda K, Woermann F, Baumgartner F, Pollmann S, Bien CG, Kissler J (2015) Investigating the brain basis of facial expression perception using multi-voxel pattern analysis. Cortex 69:131–140.

Wronka, E., & Walentowska, W. (2011). Attention modulates emotional expression processing. Psychophysiology, 48(8), 1047–1056. doi:10.1111/j.1469-8986.2011.01180.x

Zhang H, Japee S, Nolan R, Chu C, Liu N, Ungerleider LG (2016) Face-selective regions differ in their ability to classify facial expressions. Neuroimage 130:77–90.

